# Preferential phosphatidylinositol 5-phosphate binding contributes to a destabilization of the VHS domain structure of Tom1

**DOI:** 10.1101/671883

**Authors:** Wen Xiong, Tuo-Xian Tang, Evan Littleton, Arba Karcini, Iulia M. Lazar, Daniel G. S. Capelluto

**Author notes:** Correspondence and request for materials should be addressed to: Daniel G. S. Capelluto.

## Abstract

Tom1 transports endosomal ubiquitinated proteins that are targeted for degradation in the lysosomal pathway. Infection of eukaryotic cells by *Shigella flexneri* boosts oxygen consumption and promotes the synthesis of phosphatidylinositol-5-phosphate [PtdIns5P], which triggers Tom1 translocation to signaling endosomes. Removing Tom1 from its cargo trafficking function hinders protein degradation in the host and, simultaneously, enables bacterial survival. Tom1 preferentially binds PtdIns5P *via* its VHS domain, but the effects of a reducing environment as well as PtdIns5P on the domain structure and function are unknown. Thermal denaturation studies demonstrate that, under reducing conditions, the monomeric Tom1 VHS domain switches from a three-state to a two-state transition behavior. PtdIns5P reduced thermostability, interhelical contacts, and conformational compaction of Tom1 VHS, suggesting that the phosphoinositide destabilizes the protein domain. Destabilization of Tom1 VHS structure was also observed with other phospholipids. Isothermal calorimetry data analysis indicates that, unlike ubiquitin, Tom1 VHS endothermically binds to PtdIns5P through two noncooperative binding sites, with its acyl chains playing a relevant role in the interaction. Altogether, these findings provide mechanistic insights about the recognition of PtdIns5P by the VHS domain that may explain how Tom1, when in a different VHS domain conformational state, interacts with downstream effectors under *S. flexneri* infection.

## Introduction

Cells dampen signaling through a broad range of mechanisms to control cellular processes, including ubiquitinated cell-surface receptor (cargo) internalization and delivery to the surface of early endosomes ^1^. Cargo is further sorted and allocated into intralumenal vesicles in multivesicular bodies for later transportation and degradation in the lysosomal lumen. However, in some cases, ubiquitin-free receptors can be internalized in complex with their ligands to further propagate signaling from a unique population of endosomes known as signaling endosomes ^2^. Ubiquitination is a signal for protein degradation by the proteasome machinery, but it can also lead to other nondegradative pathways ^3^. Cargo sorting is mediated by ubiquitin-binding domains, which weakly bind to monomers or polymers of ubiquitin ^4^. Ubiquitin-binding domains display higher selectivity and tighter binding to certain types of polyUb chains, where these polymers adopt either packed (*i.e*., K48-linked Ub chains) or extended (*i.e*., K63-linked Ub chains) conformations ^4^.

At the endosomal surface, cargo is transported through the endosomal sorting complex required for transport (ESCRT) apparatus. The ESCRT apparatus consists of four complexes named ESCRT-0, -I, -II, and –III, which sequentially transport cargo to intralumenal vesicles where cargo is loaded ^5^. The canonical ESCRT-0 complex is composed of Hrs and STAM1/2 proteins ^6^. Both Hrs and STAM1/2 bind cargo through their double-sided Ub-interacting motif (UIM) and VHS domains ^7–9^. Anchoring of canonical ESCRT-0 to endosomal membranes is through PtdIns3P recognition via the FYVE domain of Hrs ^10^. Functionally related proteins, including the endosomal target <u>o</u>f <u>M</u>yb<u>1</u> (Tom1), and its close relatives Tom1-like 1 and 2, Toll-interacting protein, and Endofin, have been proposed to work in parallel with ESCRT-0 as alternative ESCRT-0 (alt-ESCRT-0) early cargo transporters ^11,12^. Like canonical ESCRT-0 proteins, alt-ESCRT-0 proteins have been reported to interact with ESCRT-I ^13,14^, making it possible for them to interface with the remaining ESCRT apparatus. Tom1 is a multimodular protein composed of an N-terminal VHS domain, followed by a central GAT domain, and a long C-terminal region ^15^. Both the VHS and GAT domains are involved in the recognition of cargo via their ubiquitin moieties ^8,16–18^, whereas the C-terminal region has been reported to bind to the alt-ESCRT member, Endofin ^19^.

Bacterial pathogens, including *Shigella flexneri*, *Salmonella thyphimurium*, and *Salmonella dublin*, employ metabolic strategies to alter phosphatidylinositol 5-phosphate [PtdIns5P] levels to efficiently survive with the adverse environment in the host cell ^20^. For example, *S. flexneri* invasion is thought to modify the overall organ oxygen consumption condition in the host ^21^, consequently, making a more reducing cell environment. This condition would represent the major cause of the generation of hypoxia in the host ^21^. *S. flexneri* invades host cells through internalization within a vacuole, then disrupting and escaping from it by delivering the virulent factor IpgD ^22^, a lipid phosphatase that generates endosomal PtdIns5P, using cellular phosphatidylinositol 4,5-bisphosphate ^23^. Accumulation of PtdIns5P has been reported to recruit Tom1 to signaling endosomes, thereby impairing both endosomal maturation and epidermal growth factor receptor (EGFR) degradation ^24^. Endosomal recruitment of Tom1 relies on its N-terminal VHS domain, which preferentially binds PtdIns5P ^24^. The aim of this study was to establish whether nonreducing and reducing environments and PtdIns5P binding influence the structure and, consequently, the function, of Tom1. The results identified a novel, irreversible, unfolded intermediate state in the Tom1 VHS domain under nonreducing conditions. Unlike phosphatidylcholine (PtdCho), the intermediate state of the protein was observed in the presence of PtdIns5P under both reducing and nonreducing conditions. However, PtdIns5P and related phospholipids, destabilized Tom1 VHS by reducing its thermostability and conformational compaction. Unlike ubiquitin, two PtdIns5P-binding sites, a high and a low affinity site with predominantly endothermic binding profiles, were identified in the Tom1 VHS domain with the acyl chain of the lipid playing an important role in the association. Results are discussed within the context of Tom1 recruitment to PtdIns5P-enriched endosomal compartments under bacterial infection conditions.

## Results and Discussion

### Characterization of the monomeric Tom1 VHS domain

The Tom1 VHS domain (Fig. 1a) was expressed, as a glutathione S-transferase (GST) fusion protein, in *E.coli* (Rosetta). The highest expression levels were achieved by induction with 1 mM isopropyl β-D-1-thiogalactopyranoside (IPTG) and incubation of the cells for 4 h at 25°C. Cells were lysed and the resulting cytoplasmic fraction was incubated with glutathione beads and the purified fusion protein cleaved with thrombin (Fig. 1b). Removal of protein aggregates was obtained after a second purification step using size-exclusion chromatography (data not shown). Reloading the Tom1 VHS domain in the same column, either in the absence or presence of dithiothreitol (DTT), yielded a single peak with an apparent molecular weight of 10.7 kDa (Fig. 1c and data not shown), distinct from what was predicted for a monomeric globular protein (15.7 kDa). This result suggests that Tom1 VHS is likely interacting with the size-exclusion matrix. To clarify these discrepancies, we performed infusion electrospray ionization mass spectrometry (ESI-MS) of the purified protein. A full mass spectrum of the protein is provided in Fig. 1d. The various protein charge states had a m/z representative of the Tom1 VHS domain average mass of 15,708.01, in very close agreement with the predicted molecular mass of the protein. While distinct isotope peaks for this charge state could not be produced on a low resolution linear trap quadrupole (LTQ) mass spectrometer to identify the monoisotopic peak and calculate the protein monoisotopic mass, the isotopic profile of charge state 10+ produced via an ultrazoom scan, as well as its maximum intensity isotope, corresponded to the theoretical predictions generated with the Protein Prospector program (http://prospector.ucsf.edu/prospector/cgi-bin/msform.cgi?form=msisotope). Seventeen unique tryptic products of Tom1 VHS covered ~91% of the amino acid sequence (Supplementary Table S1). Dynamic light scattering showed that 95% of Tom1 VHS was found in the peak with an intensity-average value of 4.9 nm (Fig. 1e). Overall, data indicates that the Tom1 VHS domain is intact and present in a monomeric single state.

**Figure 1:**
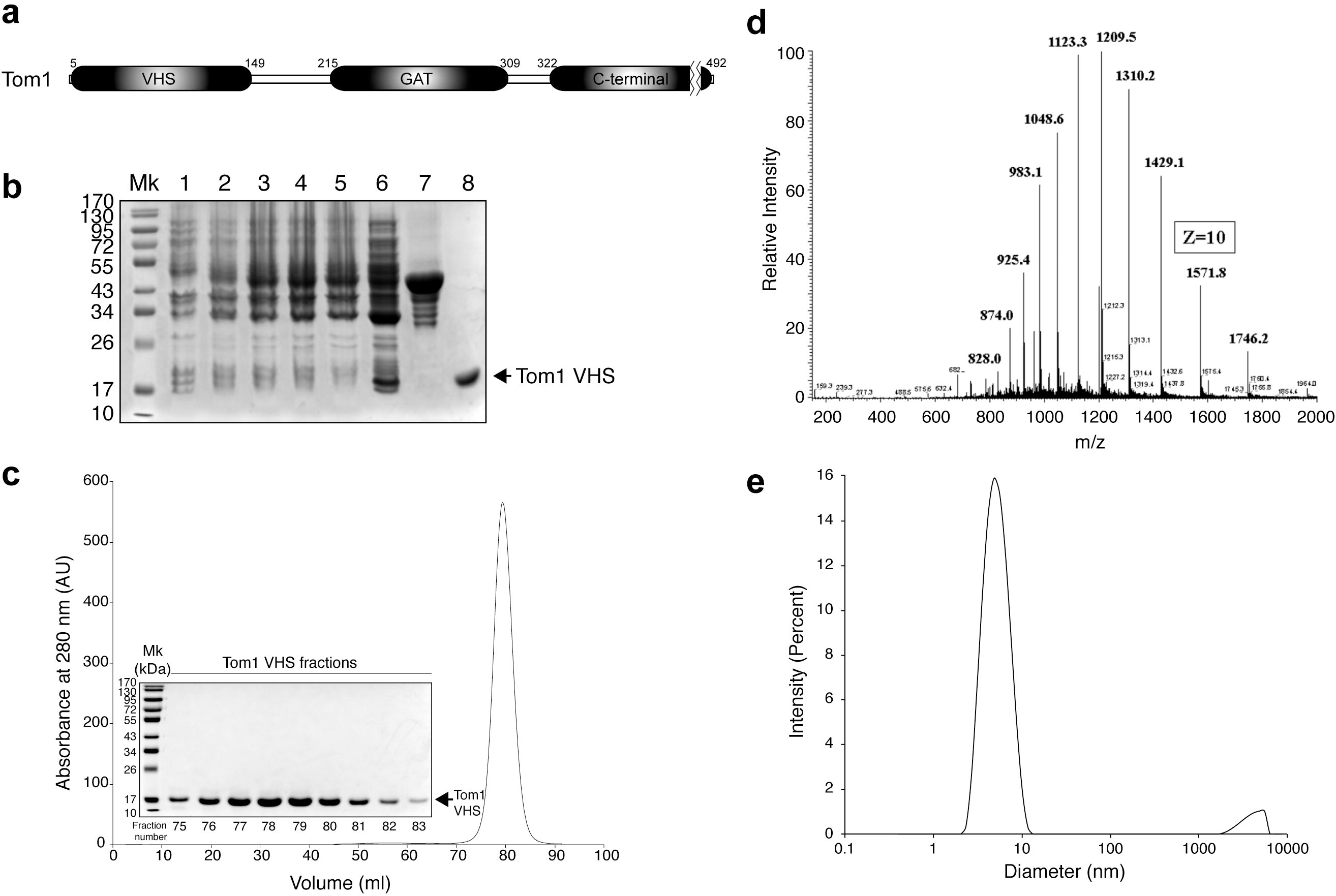
(a) Modular organization of Tom1with the domain boundaries labeled. (b) SDS-PAGE showing the expression and purification of recombinant GST-Tom1 VHS and untagged Tom1 VHS. Lane 1, cell extract before IPTG induction; lanes 2-4, cell extracts after 1, 2, and 4 h incubation with IPTG, respectively; lane 6, protein soluble cell fraction; lane 7, purified GST-Tom1 VHS bound to glutathione beads; lane 8, purified untagged Tom1 VHS. (c) Size-exclusion chromatography showing the elution of Tom1 VHS. Inset: fractions containing purified Tom1 VHS. (d) Electrospray ionization-MS analysis of recombinant Tom1 VHS. (e) DLS analysis of purified Tom1 VHS at 25°C.

### Identification of an irreversible unfolding intermediate

The Tom1 VHS domain presents three cysteine residues, one located on helix 2 (C37) and two others on helix 3 (C77 and C81) (Fig. 2a). The crystal structure of the Tom1 VHS domain was solved under reducing conditions ^25^, and therefore, the presence of a disulfide bond in the domain under physiological conditions is unknown. Both VHS domains from the Tom1-related proteins, GGA1 and GGA2, present a disulfide bond that locks the helices 2 and 4 in these proteins ^26,27^. Despite Tom1 being found at the cytosol (free or at the surface of endosomes), the locations of the cysteine residues in the VHS domain structure (Fig. 2a) are ideal for the formation of a disulfide bridge in the protein. Near-UV circular dichroism (CD) protein spectra provide information about their tertiary structure. The signal due to the excitonic coupling of the π-π* excitations on the aromatic rings of tryptophan, tyrosine, and phenylalanine could represent an extremely responsive probe for small structural changes ^28^. Under nonreducing conditions, the near-UV CD spectrum of Tom1 VHS displayed a broad minimum at 278 nm, which is associated with signals from tyrosine residues (Fig. 2b). The protein contains two tryptophan residues (W30 and W124), which do not depict a well-resolved near-UV spectral signal. The disulfide bond signal is generally found in the range of 240-290 nm, but the lack of any evident absorption in that region in the Tom1 VHS spectrum may be due to the overlapping signals from the aromatic side chains of tyrosine and phenylalanine residues. Under reducing conditions, the near-UV spectrum of Tom1 VHS showed minor changes, including a shift of the minimum to 280 nm, the presence of a tryptophan-associated shoulder at 298 nm, and some signal shifts beyond 305 nm (Fig. 2b). These results suggest that the transition from nonreducing to reducing conditions induces local conformational changes in the protein that might impact changes in its biological function.

**Figure 2:**
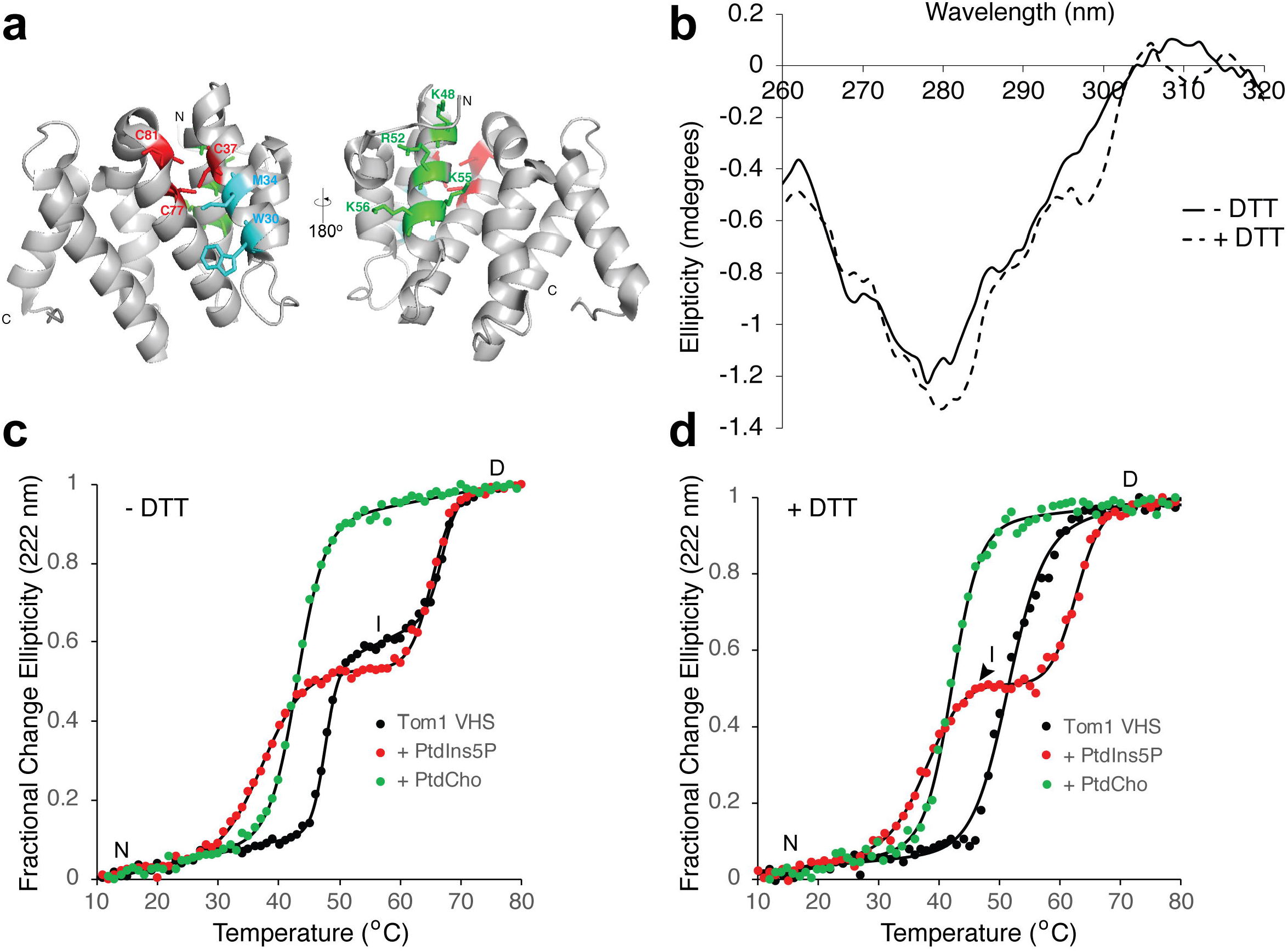
Effect of phospholipids on the thermal stability of the Tom1 VHS domain. (a) Two views of the cartoon representation of the Tom1 VHS domain indicating the location of the three conserved cysteine residues (red) and the putative PtdIns5P (green) and ubiquitin (cyan) binding sites. (b) Near-UV CD spectra of Tom1 VHS in nonreducing and reducing conditions. (c) Thermal denaturation of the Tom1 VHS domain under nonreducing conditions in the absence (black circles) and presence of a molar excess of PtdIns5P (red circles) or PtdCho (green circles). Results from one representative CD measurement of the protein are shown in the figure. Data were fitted to the merged points using a two-state denaturation model for PtdCho-bound Tom1 VHS or the three-state denaturation model for free or PtdIns5P-bound Tom1 VHS. (d) Thermal denaturation of the Tom1 VHS domain under reducing conditions in the absence (black circles) and presence of a molar excess of PtdIns5P (red circles) or PtdCho (green circles). Results from one representative CD measurement of the protein are shown in the figure. Data were fitted to the merged points using a two-state denaturation (free or PtdCho-bound Tom1 VHS) or three-state denaturation (PtdIns5P-bound Tom1 VHS) state models.

To investigate the conformational stability of the Tom1 VHS domain, we carried out thermal denaturation experiments using far-UV CD spectroscopy by looking at changes at 222 nm, where the ellipticity of the α-helix is maximal as a function of temperature under both nonreducing and reducing conditions. Under nonreducing conditions, we observed that unfolding of the native Tom1 VHS fitted well to a thermodynamic model for the presence of three-state transitions (Fig. 2c, black circles). The first transition, which led to an intermediate state of the protein, exhibited a midpoint transition of 47.5°C, whereas the second transition, which headed to a fully denatured protein, showed a melting temperature of 66.9°C (Table 1). An intermediate state, referred to as a molten globule or a non-native compact state of a protein, still exhibits secondary structure, but lacks tertiary structure due to the presence of a solvent-exposed unpacked hydrophobic core. Thermal denaturation of the Tom1 VHS domain was irreversible even at the intermediate state (Supplementary Fig. S1 and data not shown). In addition, this intermediate state exhibited poor secondary structure content as reflected by the loss of ellipticity at the far-UV spectrum of the protein measured at 55°C (Supplementary Fig. S1) with evidence of aggregation (data not shown). The loss of helical structure related with the formation of the intermediate state of Tom1 VHS was estimated to be ~77%. As occurs with the Tom1 VHS domain under nonreducing conditions, proteins that denature irreversibly mostly display a transition from the native state to the aggregated state through an unfolded intermediate state ^29^. To better understand this behavior, we calculated its hydrophobicity using the Kyte and Doolittle hydrophobicity score ^30^. Tom1 VHS displayed hydrophobicity in α-helices 2, 4, 5, 6, 7 and 8 (Supplementary Fig. S1), indicating that the intermediate unfolds irreversibly likely due to its major hydrophobic character.

**Table 1:**
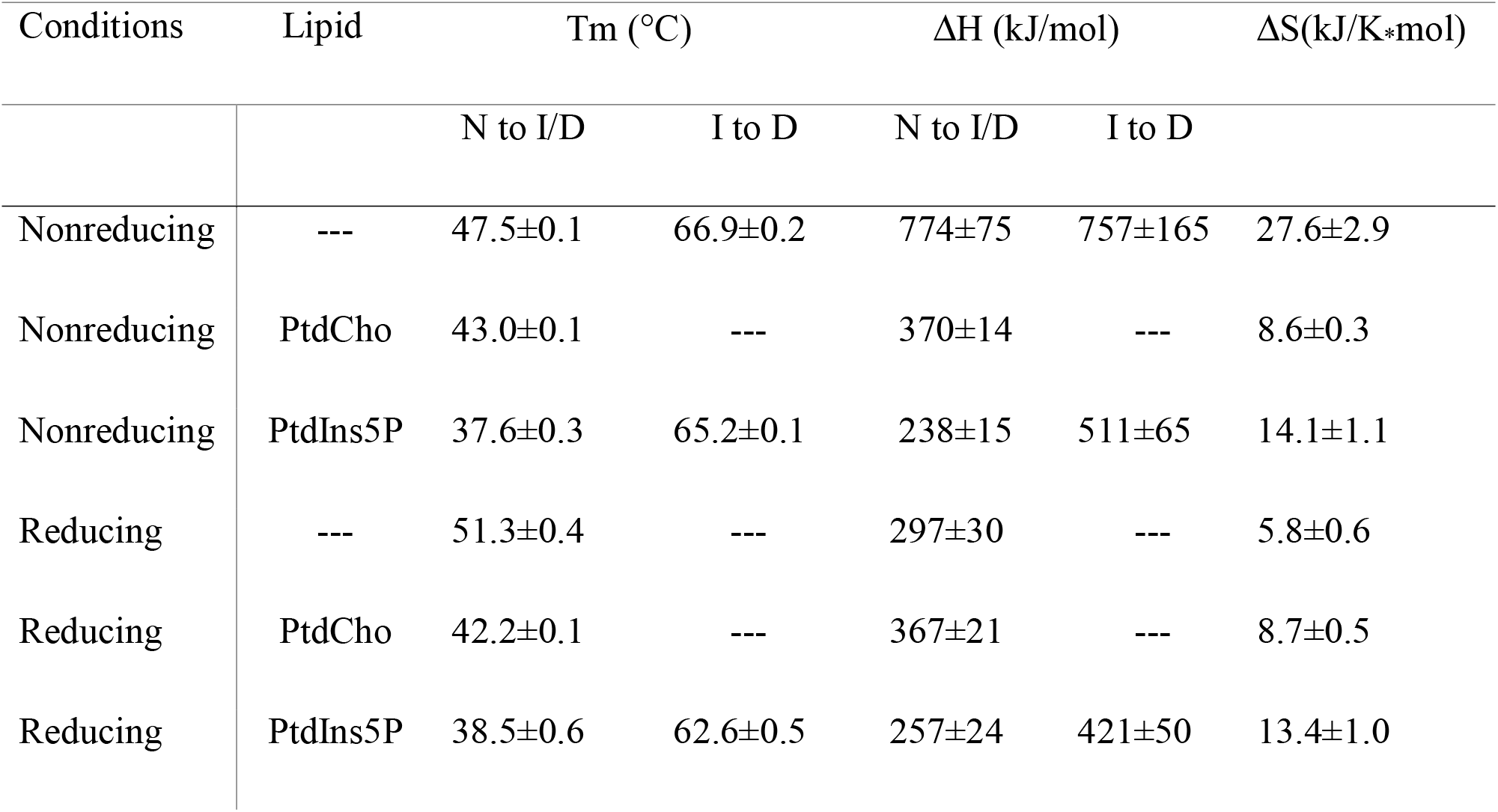
Mid-transition temperatures for Tom1 VHS under nonreducing and reducing conditions.

Consistent with previous observations ^24^, we also found that the Tom1 VHS domain binds PtdIns5P (Supplementary Fig. S2). To establish whether the interaction with PtdIns5P produces changes to the structure of the Tom1 VHS domain, the conformational stability of the protein was investigated by thermal denaturation. When Tom1 VHS was incubated with excess PtdIns5P under nonreducing conditions, the melting curve of the protein shifted down ~10°C in the native to intermediate transition, without any changes from the intermediate to denatured transition (Fig. 2c and Table 1). Therefore, association of Tom1 VHS to PtdIns5P leads to destabilization of the native state without markedly affecting the intermediate transition. Remarkably, PtdCho, the most abundant phospholipid in mammalian membranes ^31^, also destabilized Tom1 VHS but led to a two-state protein unfolding transition with a mid-transition temperature of 43°C, about 5°C higher than that in the presence of PtdIns5P (Fig. 2c and Table 1). A destabilizing effect on Tom1 VHS thermal unfolding, under nonreducing conditions, was also observed when the protein was pre-incubated with phosphatidylinositol (PtdIns) with a mid-transition temperature of 34°C but, unexpectedly, the protein remained in the intermediate state even at the highest temperature assayed (Supplementary Fig. S2). This result suggests that the presence of disulfide bonds (no DTT) in Tom1 VHS may impair the protein to fully denature when PtdIns is present in excess. Nonetheless, overall data indicate that phospholipids can be recognized by the Tom1 VHS domain, albeit with different avidities ^24^, and their excess cause destabilizing effects on the protein thermal unfolding.

Interestingly, under reducing conditions, where the intramolecular disulfide bonds are lost, the Tom1 VHS displayed a distinct cooperative two-state transition with a mid-transition temperature of 51.3°C (Fig. 2d). These results suggest that a disulfide bridge, which can form close to the N-terminus of the protein (Fig. 2a) under nonreducing conditions, impairs cooperative unfolding behavior (Fig. 2c). In the presence of PtdIns5P and under reducing conditions, however, Tom1 VHS retains the noncooperative three-state unfolding transition with the intermediate state being less stable than that observed under nonreducing conditions (Fig. 2c-d). Interestingly, PtdIns switched the thermal denaturation of Tom1 VHS from a two to a three-state fully denatured transition, under reducing conditions (Supplementary Fig. S2). On the other hand, Tom1 VHS still displayed a two-transition unfolding pattern in the presence of PtdCho (Fig. 2d). Unfolding of Tom1 VHS was irreversible under reducing conditions as well (data not shown), thus, excluding the possibility that the irreversibility of unfolding is due to the presence of a disulfide bridge.

To further investigate the PtdIns5P-dependent destabilizing effects in the Tom1 VHS structure, we analyzed its susceptibility to trypsin cleavage. Whereas Tom1 VHS was proteolyzed at high trypsin concentrations, addition of PtdIns5P triggered heavy digestion of the protein even at the lowest trypsin amount tested (500 ng) (Fig. 3a and Supplementary Fig. S3). As Tom1 VHS exhibits broad preference for phosphoinositides ^24^, the protein also showed heavy proteolysis when pre-incubated with an excess of either PtdIns or PtdIns3P (Supplementary Fig. S3 and S4). Addition of an excess of PtdCho also led to more sensitivity of Tom1 VHS to trypsin, but proteolysis was less evident than that observed in the presence of phosphoinositides or PtdIns (Fig. 3a and Supplementary Fig. S3). In contrast, the Tom1 VHS binding partner, ubiquitin, did not destabilize the protein and mirrored the trypsin digestion of free Tom1 VHS (Fig. 3a and Supplementary Fig. S3). Taken together, these results indicate that binding of the Tom1 VHS domain to an excess of PtdIns and phosphoinositides, and perhaps PtdCho to a much lesser extent, induces major tertiary structural changes in the protein that lead to a loss of its conformational compaction and, consequently, less thermostability and more susceptibility to limited trypsin proteolysis. Given that Tom1 VHS has a preference for PtdIns5P ^24^, we speculate that these conformational changes, driven by the phosphoinositide, might favor the contact of Tom1 with other macromolecules that are required to transduce PtdIns5P-dependent signaling events. In this context, the PtdIns5P-dependent recruitment of Tom1 at signaling endosomes inhibits endosomal maturation and EGFR degradation, with a consequential enhancement of pathogen survival ^24^. Although Tom1 does not serve as an alt-ESCRT-0 protein for EGFR at signaling endosomes ^24^, it is still unknown how Tom1 facilitates the intracellular accumulation of EGFR under bacterial infection. Interestingly, the endosomal intercellular adhesion molecule-1 is degraded by the host lysosomal pathway in a bacterially-produced PtdIns5P-dependent manner, but the role of Tom1 in this process has not yet been addressed ^32^.

**Figure 3:**
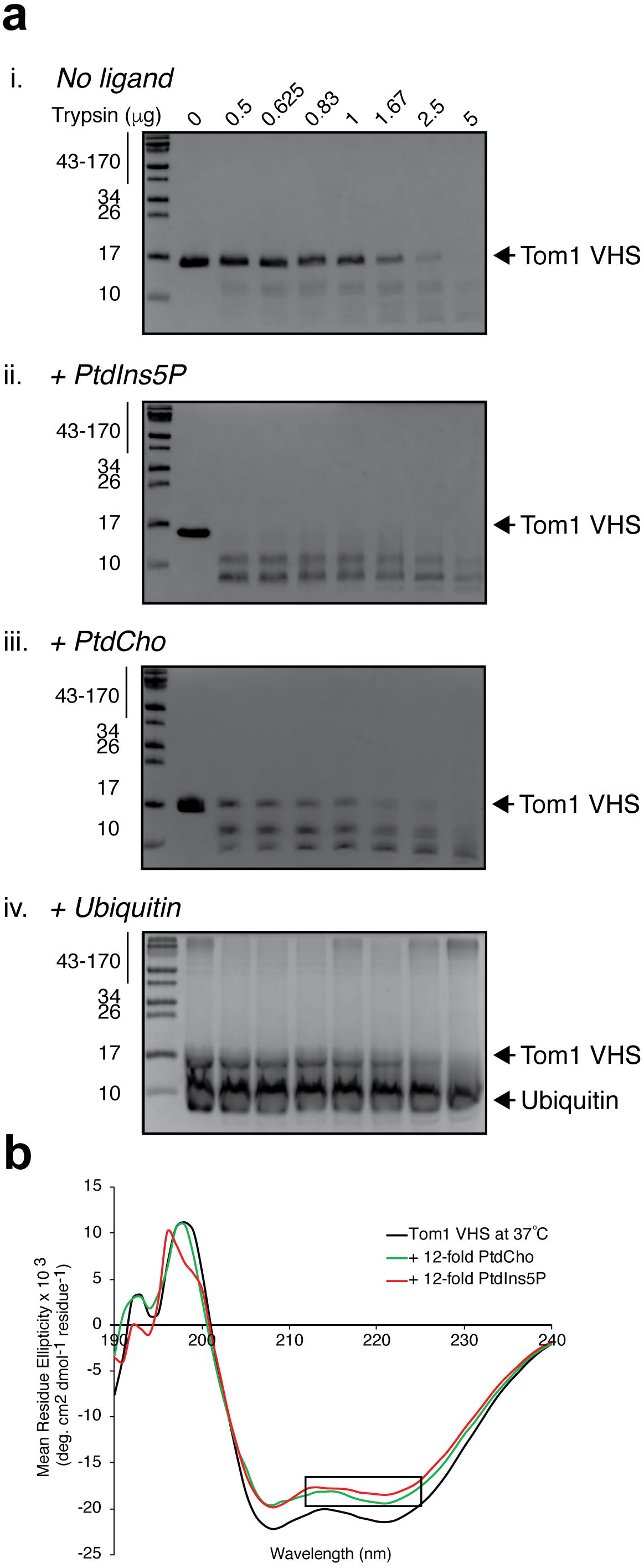
PtdIns5P-dependent destabilization of the Tom1 VHS domain. (a) SDS-PAGE showing the trypsin limited proteolysis of Tom1 VHS in the absence (i) and presence of PtdIns5P (ii), PtdCho (iii), and ubiquitin (iv). (b) Far-UV CD spectra of Tom1 VHS in the absence (black) and presence of PtdIns5P (red) or PtdCho (green) under nonreducing conditions at 37°C. The loss of signal at 218-222 nm is boxed.

### Structural features of Tom1 VHS upon PtdIns5P binding under nonreducing and reducing conditions

We next examined the secondary structural changes by tracking the ellipticity of the protein in the far-UV CD region under nonreducing conditions. At 25°C, Tom1 VHS exhibited two minima at 208 and 222 nm with an estimated 67% of helical structure (Supplementary Fig. S5 and Supplementary Table S2), an identical value deduced from the crystal structure of the protein ^25^. Extensive interhelical contacts in proteins can be predicted by estimating their 222/208 ellipticity ratio. A value of ~1.0 suggests that α-helices display extensive interhelical contacts in helical bundles and coiled coil regions, whereas a value of ~0.8 indicates limited interhelical contacts ^33^. The 222/208 ellipticity ratio value for Tom1 VHS at 25°C was 0.976±0.004. Consequently, our structural analysis indicates that there are extensive interhelical contacts in the structure of the Tom1 VHS domain under nonreducing conditions. The addition of either PtdCho or PtdIns5P did not markedly change the 222/208 ratio, with values of 0.968±0.008 and 0.970±0.000, respectively.

At nearly physiological conditions (37°C), the far-UV CD spectrum of the protein slightly decreased in ellipticity with an estimated helical content of 62% and a 222/208 ratio of 0.957±0.004 (Fig. 3b and Table 2). Unlike what is observed at 25°C (Supplementary Fig. S5 and Supplementary Table S2), incubation of Tom1 VHS with an excess of soluble PtdIns5P at 37°C and under nonreducing conditions, led to a loss of CD ellipticity with an evident decrease in the 208-222 nm band (mostly in the 218-222 nm region) and with a 222/208 ellipticity ratio of 0.932±0.011, suggesting that PtdIns5P reduces interhelical contacts (Fig. 3b). In the presence of PtdCho, a decrease in the 208-222 nm ellipticity band was also observed but without a further loss in the 218-222 nm band (boxed region, Fig. 3b), giving a 222/208 ellipticity ratio of 0.972±0.010. Similar findings were observed when the CD spectra of Tom1 VHS were collected under reducing conditions (Supplementary Fig. S5; Table 2 and Supplementary Table S2). All together, these results show that PtdIns5P destabilizes Tom1 VHS independent of a reducing environment.

**Table 2:** Estimation of the secondary structure content of Tom1 VHS under nonreducing and reducing conditions and at 37°C using the CDSSTR algorithm.

| Lipid | DTT | Content (%) |  |  | Random coil | NRMSD |
| --- | --- | --- | --- | --- | --- | --- |
| | | α-helix | β-strand | Turns | | |
| - | - | 62 | 12 | 10 | 16 | 0.004 |
| PtdCho | - | 59 | 14 | 11 | 16 | 0.004 |
| PtdIns5P | - | 59 | 14 | 10 | 17 | 0.004 |
| - | + | 63 | 17 | 13 | 7 | 0.003 |
| PtdCho | + | 56 | 18 | 13 | 13 | 0.003 |
| PtdIns5P | + | 55 | 14 | 10 | 21 | 0.005 |

The two tryptophan residues of the Tom1 VHS domain can serve as intrinsic probes for tracking interactions with protein ligands. Under nonreducing conditions, the protein exhibited a maximum tryptophan fluorescence emission at 343 nm. Gradual addition of PtdIns5P led to a reduction in fluorescence intensity of the Tom1 VHS domain without any evident change in the fluorescence spectrum maximum (Fig. 4a) but with a biphasic-like pattern (Inset, Fig. 4a), reminiscent of two-binding events. Quenching in the Tom1 VHS emission spectra might be related to local conformational transition upon PtdIns5P binding, which can affect the local environment of one or both tryptophan side chains of the protein. PtdIns3P, PtdIns, and PtdCho also quenched the fluorescence signal, to a lesser extent, and displayed a close to saturation pattern (Supplementary Fig. S6). Under reducing conditions, PtdIns5P, PtdIns3P, and PtdIns (the last two to a slightly lesser extent) also quenched the fluorescence emission of Tom1 VHS with a biphasic-like pattern (Supplementary Fig. S6).

**Figure 4:**
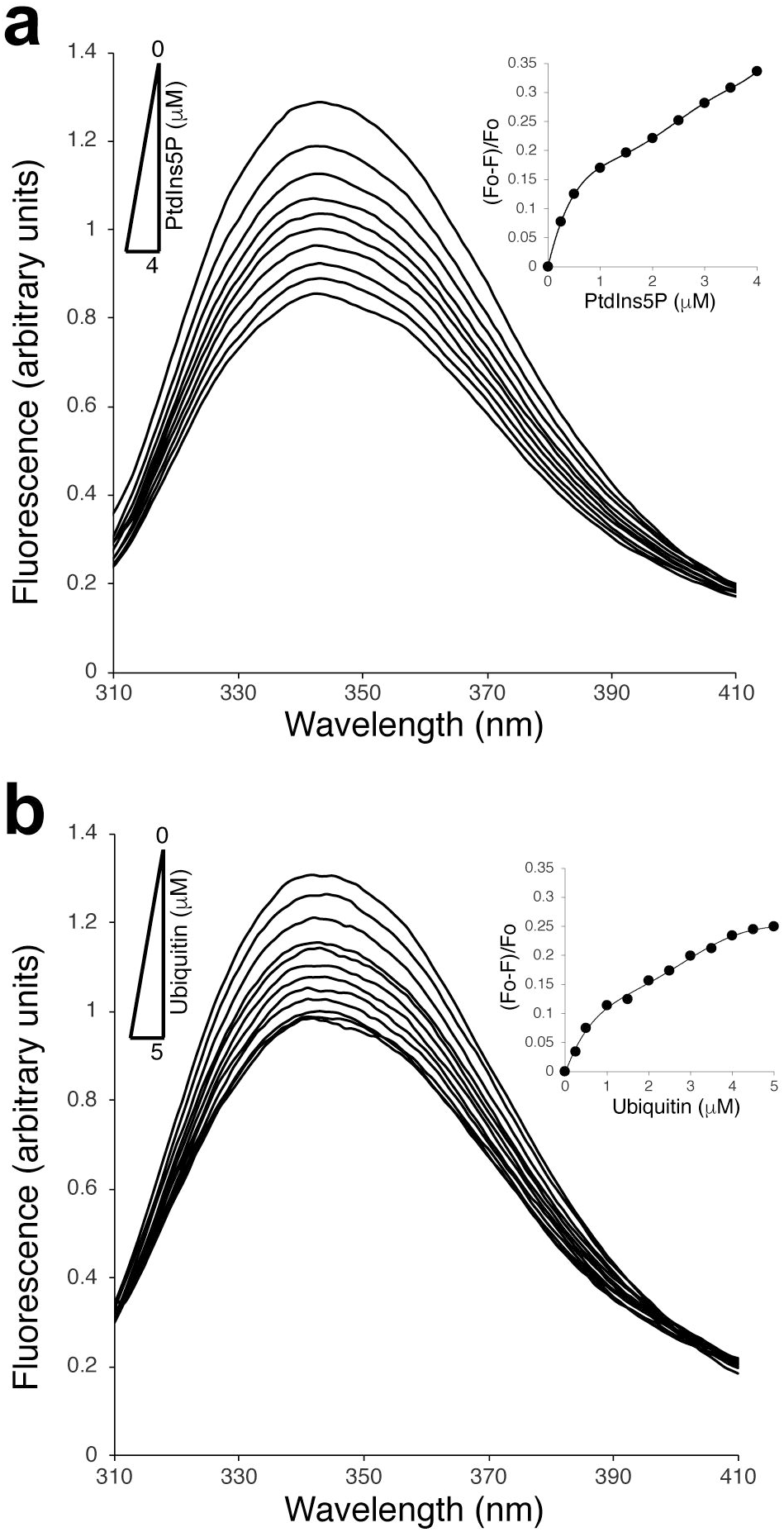
Conformational changes of Tom1 VHS associated to PtdIns5P and ubiquitin binding under nonreducing conditions. (a) Intrinsic tryptophan fluorescence of Tom1 VHS in the absence and presence of increasing concentrations of PtdIns5P. Inset: Extent of PtdIns5P-dependent quenching of Tom1 VHS signal. (b) Intrinsic tryptophan fluorescence of Tom1 VHS in the absence and presence of increasing concentrations of ubiquitin. Inset: Extent of ubiquitin-dependent quenching of Tom1 VHS signal.

Most of the VHS domains have been shown to associate to mono- and poly-ubiquitin chains covalently bound to cargo receptors. Details of the molecular association comes from structural studies for the formation of the STAM1 VHS-ubiquitin complex ^8^. STAM1 VHS recognizes the canonical hydrophobic patch at the surface of ubiquitin, represented by residues I44 and G47, which are completely buried at the complex interface ^8^. Ubiquitin recognition is mediated by the α-helices α2 and α4 with W26 and L30 playing a major role in the interaction ^8,34^. The equivalent residues in the Tom1 VHS domain are W30 and M34 (cyan residues, Fig. 2a). Ubiquitin quenched the fluorescence emission of Tom1 VHS in a saturable manner (Fig. 4b). Thus, the observed ubiquitin-mediated reduction of the fluorescence signal of Tom1 VHS (Fig. 4b) can be attributed at changes in the local environment around the indole ring of W30 as ubiquitin lacks tryptophan residues.

### Thermodynamics of Tom1 VHS association to PtdIns5P monitored by isothermal calorimetry

To better understand the PtdIns5P binding mechanism of Tom1 VHS, we carried out isothermal calorimetry (ITC) experiments. Under nonreducing conditions, binding of water soluble PtdIns5P to Tom1 VHS was markedly endothermic with the resulting isotherms fitting a two-site binding model (Fig. 5a), in line with the observed behavior of the protein from tryptophan fluorescence analysis (Fig. 4a and Supplementary Fig. S6). Tom1 VHS bound PtdIns5P with dissociation constants, *K*_D1_ and *K*_D2_, of 3.1 and 110 μM, respectively (Table 3), suggesting the presence of high and low affinity PtdIns5P binding sites in the protein. Despite the critical micelle concentration (CMC) of PtdIns5P (dioctanoyl form) is still unknown, the CMC of dioctanoyl PtdIns3P is 700±200 μM ^35^; thus, it is conceivable to assess the CMC of dioctanoyl PtdIns5P to be ~700 μM. Considering that the final concentration of PtdIns5P in the ITC sample cell ranges between 60-600 μM during a titration, most of the lipid is expected to be predominantly in a monodispersed form. It is still possible that Tom1 VHS can interact with micellar PtdIns5P, which may be responsible for the second binding event. We, therefore, evaluated the association of Tom1 VHS with the PtdIns5P headgroup, inositol 1,5 bisphosphate [Ins(1,5)P_2_], which is unable to form micelles. Binding of Tom1 VHS to Ins(1,5)P_2_ remained endothermic and best fit a single-site binding model, but the affinity for the headgroup was weak with a *K*_D_ of 213 μM (Fig. 5b, Table 3). Thus, the PtdIns5P acyl chains markedly contribute to the strength of binding to the Tom1 VHS domain. Similarly, stable association of HIV-1 Gag to membrane phosphatidylinositol 4,5-bisphosphate requires the presence of specific acyl chains in the lipid ^36^. Likewise, TIRAP requires both the headgroup and acyl chains of phosphatidylinositol 4,5-bisphosphate for its recognition ^37^. A positively charged region in Tom1 VHS, including the α3 residues K48, R52, K55, and K56 (green residues, Fig. 2a), was proposed to be the single PtdIns5P binding site ^24^. This conclusion, however, was reached based on alanine mutagenesis of these residues without any evidence of whether the protein retained proper folding upon the introduction of these mutations. The presence of two PtdIns3P/PtdIns(3,5)P_2_-binding sites has also been identified with atomic details in the yeast β-propeller PROPPIN protein, Hsv2 ^38,39^. Further studies demonstrated that the PROPPIN family members Atg18, Atg21, and Hsv2 require both PtdIns3P/PtdIns(3,5)P_2_-binding sites and an insertion loop, all located within a single domain, for membrane binding ^40^. In some cases, such as for Atg18, phosphoinositides promote protein oligomerization and accommodate the oligomer perpendicularly to the membrane in a slightly tilted orientation ^41^. Alternatively, two distinct phospholipid-binding sites are found in single protein domains such as PH, PX, and C2 domains ^42,43^. This mechanism may serve as two independent points of membrane contacts, permitting coincidence detection. Thus, the low affinity PtdIns5P binding site in Tom1 VHS might represent a site for weak binding for the phosphoinositide or for another phospholipid. It is also possible that the second binding event constitutes a nonspecific PtdIns5P-binding site in the protein at excess of lipid. Nevertheless, additional molecular studies are needed to identify the PtdIns5P binding site/s in the Tom1 VHS domain. Our ITC analysis shows positive ∆H (enthalpy) values for the association of Tom1 VHS to PtdIns5P (Table 3). The presence of PtdIns5P likely disrupts energetically favorable interactions between the atoms and water molecules in the lipid binding pocket/s of Tom1 VHS. An unfavorable enthalpy input (ΔH > 0) should require a large favorable entropy change (ΔS > 0) to have a negative free energy of association (ΔG < 0). Therefore, unfavorable ∆H contributions during the PtdIns5P binding to Tom1 VHS are compensated by an entropy-driven process (Table 3). Both far-UV CD and fluorescence data analysis suggest that PtdIns5P promotes conformational changes, including reduction of the helical content, and probably partial desolvation in the protein that might contribute to the observed entropy in this process. Unexpectedly, the protein poorly interacted with PtdCho using the same experimental conditions (Fig. 5c). We speculate that association of Tom1 VHS to PtdCho is purely entropic as this property does not give rise to a heat signal ^44^. Under the same ITC experimental conditions, we found that binding of ubiquitin to Tom1 VHS was enthalpically-driven and data best fit a one-site binding model with a *K*_D_ of 695 μM (Fig. 5d and Table 3), a value within the range reported by Ren and Hurley for a variety of VHS domains ^8^.

**Table 3:** Thermodynamic parameters of PtdIns5P, Ins(1,5)P_2_, and ubiquitin binding to the Tom1 VHS domain. Values represent the mean of at least two independent experiments. Errors are displayed as standard deviation values.

| Ligand | Mode | $K_D$ (μM) | $\Delta H$<br>(kcal. mol<sup>-1</sup>) | T $\Delta S$<br>(kcal. mol<sup>-1</sup>) | $\Delta G$<br>(kcal. mol<sup>-1</sup>) |
| --- | --- | --- | --- | --- | --- |
| PtdIns5P | High affinity | 3.1±0.6 | 0.29±0.06 | 7.81±0.05 | -7.53±0.11 |
| | Low affinity | 110.0±26.8 | 1.94±0.30 | 7.33±0.28 | -5.41±0.14 |
| Ins(1,5)P<sub>2</sub> | Single site | 213.5±67.2 | 0.88±0.35 | 5.91±0.16 | -5.03±0.19 |
| Ubiquitin | Single site | 695.0±134.0 | -6.16±0.45 | 1.84±0.33 | -4.32±0.12 |

**Figure 5:**
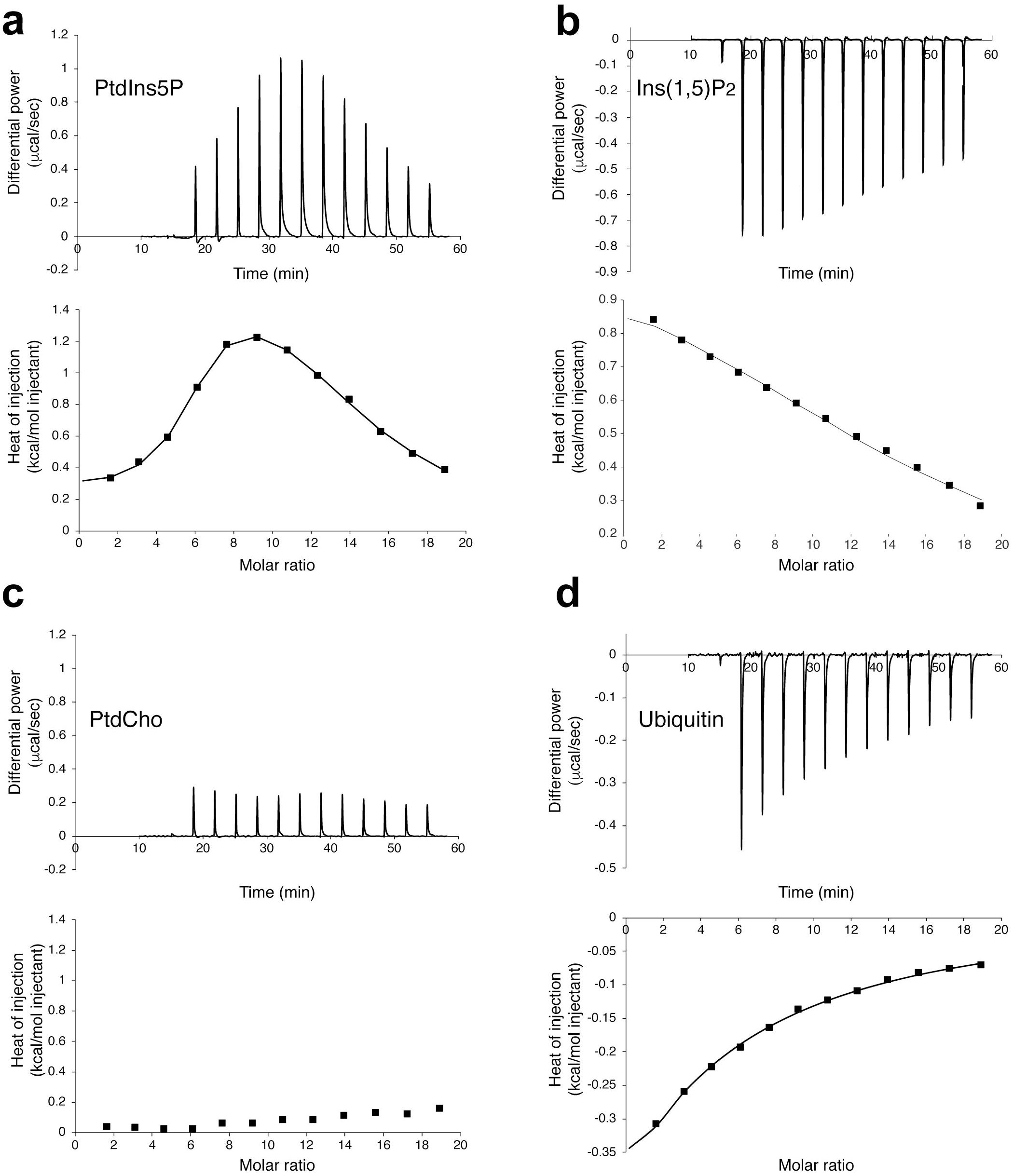
Tom1 VHS calorimetric titration curves for the association to PtdIns5P (a), Ins(1,5)P_2_ (b), PtdCho (c), and ubiquitin (d). The upper panels depict the heat change produced upon successive injections of the indicated ligand to the Tom1 VHS domain. The lower panels display the integrated binding isotherms as a function of ligand/protein ratio. Binding isotherms were fitted to the subtracted data using two-sites (PtdIns5P) or one-site (Ins(1,5)P_2_ and ubiquitin) binding models. Each titration was carried out at least two times and representative results are shown.

## Conclusions

The structural and thermodynamic results shown here provide new insights into phosphoinositide-binding domains and their behavior under nonreducing and reducing conditions. Specifically, thermal unfolding studies of the monomeric Tom1 VHS domain under nonreducing conditions show that the protein transitions through an intermediate state, which is absent under reducing conditions. Binding of Tom1 VHS to PtdIns5P leads to local conformational changes that reduce thermostability, interhelical contacts, and conformational compaction in the protein. Thus, conformation of the PtdIns5P-bound Tom1 VHS, found at a physiological temperature, may have a role in the progress of bacterial infection by facilitating yet-to-be-determined molecular interactions. In addition, the PtdIns5P-dependent conformational change of the VHS domain may explain why Tom1 is no longer engaged in ubiquitinated protein trafficking under bacterial infections. Our studies indicate that other phospholipids, including PtdIns3P, PtdIns, and PtdCho, also interact to and destabilize Tom1 VHS. Further, thermodynamic analysis suggests that Tom1 VHS displays two, both a high and a low, binding sites for PtdIns5P. Unlike ubiquitin binding, PtdIns5P binding events are markedly entropy-driven, noncooperative, and display distinct endothermic processes. Interestingly, the acyl chains of the phosphoinositide exhibit a relevant role for recognition, results that can explain the broad specificity of Tom1 VHS for phospholipids. Future studies are needed to understand the molecular mechanism by which Tom1 and its VHS domain associate to PtdIns5P-enriched membranes and to identify what regions in VHS undergo major conformational changes that trigger destabilization.

## Methods

### Protein expression and purification

The human Tom1 VHS domain was cloned as previously reported ^45^. A construct expressing human ubiquitin, cloned into the pET24d vector, was a gift from Dr. Julie Forman-Kay (University of Toronto). Ubiquitin was expressed and purified as described ^46^.

### Size-exclusion chromatography

The purified Tom1 VHS domain was loaded onto an FPLC system using a Superdex 75 column (GE Healthcare) equilibrated with 50 mM Tris-HCl (pH 7.3), and 500 mM NaCl in the absence and presence of 1 mM DTT. Protein standards for the Superdex 75 column were as follows: bovine serum albumin (66 kDa; 3.5 Å), ovalbumin (43 kDa; 2.8 Å), carbonic anhydrase (29 kDa; 2.36 Å), chymotrypsin A (25 kDa; 2.17 Å), cytochrome C (12.5 kDa; 1.7 Å), aprotinin (6.5 kDa; 1.35 Å), and vitamin B12 (1.35 kDa; 0.85 Å).

### Infusion electrospray ionization mass spectrometry and liquid chromatography-tandem mass analyses

Tom1 VHS was desalted with SPEC-PTC18 cartridges (Agilent Technologies, Santa Clara, CA) and prepared for mass spectrometry (MS) analysis in a solution of CH_3_CN/H_2_O/CH_3_COOH (50:50:1, v/v) at 10 µM concentration. The protein solution was delivered with the aid of a syringe pump (Harvard Apparatus, Holliston, MA) at 0.5 µL/min to the ion source of an LTQ mass spectrometer (Thermo Fisher Scientific, San Jose, CA). The sample was electrosprayed at 2.2 kV, and data acquisition was performed by averaging five scans per mass spectrum.

### Dynamic light scattering

DLS measurements were recorded at 25°C using a Malvern Zetasizer Nano-ZS instrument. Tom1 VHS (0.8 mg/ml) was prepared in 20 mM HEPES (pH 7.4) and 150 mM KCl. Each measurement was collected for 120 s and three accumulated spectra were averaged with the protein previously equilibrated for 2 min. Data was analyzed using the Malvern Zetasizer software v. 7.11.

### Circular dichroism

Far-UV CD measurements were acquired on a Jasco J-815 spectropolarimeter with a temperature-controlled cell holder connected to a Peltier unit at the indicated temperatures. Tom1 VHS (20 μM) was prepared in 5 mM sodium citrate (pH 7.3) and 50 mM KF with and without 0.1-5 mM DTT. Three accumulated CD protein spectra, in the absence and presence of either dioctanoyl PtdIns5P (Echelon Biosciences) or dioctanoyl PtdCho (Avanti Lipids), were recorded in a 1-mm path length quartz cell using a bandwidth of 1-nm and a response time of 1 s at a scan speed of 50 nm/min. Secondary structure content of the Tom1 VHS domain, under different experimental conditions, was estimated using the CDSSTR algorithm available at the DICHROWEB server ^47^. All CD protein spectra were corrected for buffer background. Near-UV measurements were performed in 1 mm cells containing 200 μM Tom1 VHS in 5 mM sodium citrate (pH 7.3) and 50 mM KF with and without 1 mM DTT. A scan of buffer was subtracted from the corresponding averaged protein spectra. Temperature-dependent unfolding of Tom1 VHS was followed by changes of the CD signals at 222 nm as a function of increasing the temperature from 10 to 80°C, with 1°C steps and with a temperature gradient of 0.75 °C/min. All CD measurements were recorded in duplicate. To calculate the thermodynamic parameters, the derivatives of ellipticity as a function of temperature were calculated and fitted using CDpal ^48^ following either two-or three-state denaturation models, as indicated. Thermal denaturation data were normalized between 0 and 1, with 1 denoting the unfolded protein value.

### Tryptophan fluorescence

Intrinsic tryptophan fluorescence measurements were collected using a Jasco J-815 spectropolarimeter. Emission spectra of the Tom1 VHS domain (0.25 μM) were recorded in 5 mM sodium citrate (pH 7.3) and 50 mM KF without or with 1.25 μM DTT, after excitation of the protein at 295 nm. Protein emission spectra, in the absence and presence of PtdIns5P, PtdIns3P, PtdIns, PtdCho, or ubiquitin, were collected between 310-410 nm using a 10-mm quartz cuvette at 25°C.

### Limited proteolysis

Limited proteolysis with trypsin was carried out using 10 μg of purified Tom1 VHS in the absence or presence of either PtdIns5P, PtdIns3P, PtdIns, PtdCho, or ubiquitin (1:12 molar ratio). The mixtures were incubated for 5 min at room temperature. Then, trypsin (MP Biochemicals) was added to the mixtures at amounts ranging from 0 to 5 μg for 30 min at 37°C. The reactions were quenched by the addition of SDS-PAGE loading buffer and analyzed by 10% Tricine-SDS-PAGE and stained with Coomassie Brilliant Blue G-250 (VWR).

### Isothermal titration calorimetry

ITC experiments were performed in 20 mM HEPES (pH 7.4) and 150 mM KCl. Proteins and lipids were dissolved in this buffer. Binding traces were analyzed with a MicroCal PEAQ (Malvern) equilibrated at 25°C. PtdIns5P (5 mM), PtdCho (5 mM), or ubiquitin (5 mM) were placed in the syringe, whereas the Tom1 VHS concentration in the sample cell for every titration was 50 μM. Titrations were carried out with 12-fold 3 μl injections. The first injection, which was 0.4 μl, was not considered for data analysis. All experiments were carried out at least in duplicate. The reported heats of binding were established by integrating the experimental peaks after adjusting the effect of heat dilution of either the lipids, Ins(1,5)P_2_, or ubiquitin into buffer in a control measurement. The integrated heat data were fit with the one-(ubiquitin and Ins(1,5)P_2_) or two-binding (PtdIns5P) site models using the MicroCal PEAQ-ITC analysis software.

## Acknowledgments

We thank Dr. Janet Webster for critical reading on the manuscript. This research was supported in part by the Virginia Academy of Sciences (to E.L.).

## Author Contributions

W.X., I.M.L., and D.G.S.C. conceived the experiments. W.X., TX.T., and E.L. purified proteins and performed lipid-protein overlay assays. W.X. performed and analyzed CD, tryptophan fluorescence, and ITC experiments. TX.T. performed and analyzed dynamic light scattering and trypsin limited proteolysis experiments. A.K. and I.M.L. carried out the mass spectrometry experiments and analysis. D.G.S.C. wrote the manuscript with all authors contributing to the discussion and revision of the manuscript.

The datasets generated during and/or analyzed during the current study are available from the corresponding author by reasonable request.

