## Supplementary Material for "Preferential phosphatidylinositol 5-phosphate binding contributes to a destabilization of the VHS domain structure of Tom1"

**Methods**

**Trypsin proteolysis and mass spectrometry analysis.** The sequence of the protein was verified by performing liquid chromatography (LC)-tandem MS analysis of the protein digest. For this purpose, the protein was denatured with urea (8 M) and reduced with DTT (5 mM) for 1 h at 55-57ºC. After 10-fold dilution with $NH_4HCO_3$ (50 mM) to a concentration of 0.5 μM, protein was subjected to proteolysis with MS sequencing grade trypsin (Promega, Madison, WI) overnight, at 37°C (substrate:trypsin 50:1 w/w). After cleanup with a SPEC-PTC18 cartridge, the sample re-dissolved in a solution of $CH_3CN/H_2O$/TFA (2:98:0.01, v/v) and analyzed by LC-MS with a micro-HPLC system (Agilent Technologies 1160 micro-LC) adapted in-house with a nano-ESI source,[1] and LC eluent delivered at ~180 nL/min and ESI voltage set to 2.2 kV. The 4 h long gradient involved increasing the concentration of solvent B ($H_2O$:$CH_3CN$:TFA, 10/90/0.01 v/v) from 10 % to 100 %. The composition of solvent A was $H_2O$:$CH_3CN$:TFA, 96/4/0.01 v/v. Only the top 5 most intense peaks from each mass spectrum were considered for data dependent tandem MS analysis[2]. The database search was completed by using a UniProt *E.coli* database appended with the amino acid sequence of the protein, and the Proteome Discoverer 2.3 (Thermo Electron) software package. Two missed cleavages were allowed in the search, and the false discovery rate was set at <1 %[2].

**Lipid-protein overlay assay.** PtdIns(5)P strips were prepared by immobilizing 1 μl of the indicated amount of the phosphoinositide solubilized in chloroform/methanol/water (65:35:8) onto Protran nitrocellulose membranes (GE Healthcare). PtdIns(5)P-bound strips were blocked overnight with 3% fatty acid-free BSA (Sigma) in 20 mM Tris-HCl (pH 8.0), 150 mM NaCl, and 0.1 % Tween-20 at 4°C. Then, strips were incubated with proteins (60 nM) in the same buffer for 1 h at 4°C. Strips were washed four times with the same buffer followed by incubation with a rabbit anti-glutathione S-transferase antibody (Proteintech). Following an additional washing step with the same buffer, strips were incubated with a donkey anti-rabbit-horseradish peroxidase antibody (GE Healthcare). The presence of the PtdIns(5)P-bound protein was probed using Supersignal West Pico chemiluminescent reagent (Pierce).

**Supplementary Table S1.** Identification of the Tom1 VHS domain. MS/MS results obtained on peptides generated by trypsin proteolysis of Tom1 VHS.

| Peptide | Tom1 | Sequence |
|---|---|---|
| 1 | 17-52 | IEKATDGSLQSEDWALNMEICDIINETEEGPKDALR |
| 2 | 20-48 | ATDGSLQSEDWALNMEICDIINETEEGPK |
| 3 | 20-52 | ATDGSLQSEDWALNMEICDIINETEEGPKDALR |
| 4 | 20-55 | ATDGSLQSEDWALNMEICDIINETEEGPKDALRAVK |
| 5 | 57-79 | RIVGNKNFHEVMLALTVLETCVK |
| 6 | 58-79 | IVGNKNFHEVMLALTVLETCVK |
| 7 | 58-84 | IVGNKNFHEVMLALTVLETCVKNCGHR |
| 8 | 63-79 | NFHEVMLALTVLETCVK |
| 9 | 63-101 | NFHEVMLALTVLETCVKNCGHRFHVLVASQDFVESVLVR |
| 10 | 80-101 | NCGHRFHVLVASQDFVESVLVR |
| 11 | 85-101 | FHVLVASQDFVESVLVR |
| 12 | 102-116 | TILPKNNPPTIVHDK |
| 13 | 102-129 | TILPKNNPPTIVHDKVLNLIQSWADAFR |
| 14 | 107-129 | NNPPTIVHDKVLNLIQSWADAFR |
| 15 | 117-129 | VLNLIQSWADAFR |
| 16 | 117-146 | VLNLIQSWADAFRSSPDLTGVVTIYEDLRR |
| 17 | 130-145 | SSPDLTGVVTIYEDLR |

**Supplementary Table S2:**

Estimation of the secondary structure content of Tom1 VHS under nonreducing and reducing conditions at 25°C using the CDSSTR algorithm.

| | | Content (%) | | | | |
|---|---|---|---|---|---|---|
| Lipid | DTT | α-helix | β-strand | Turns | Random coil | NRMSD |
| - | - | 67 | 10 | 10 | 13 | 0.003 |
| PtdCho | - | 66 | 13 | 9 | 12 | 0.004 |
| PtdIns5P | - | 66 | 14 | 9 | 11 | 0.003 |
| - | + | 65 | 15 | 9 | 11 | 0.004 |
| PtdCho | + | 65 | 16 | 8 | 11 | 0.003 |
| PtdIns5P | + | 62 | 19 | 10 | 9 | 0.003 |

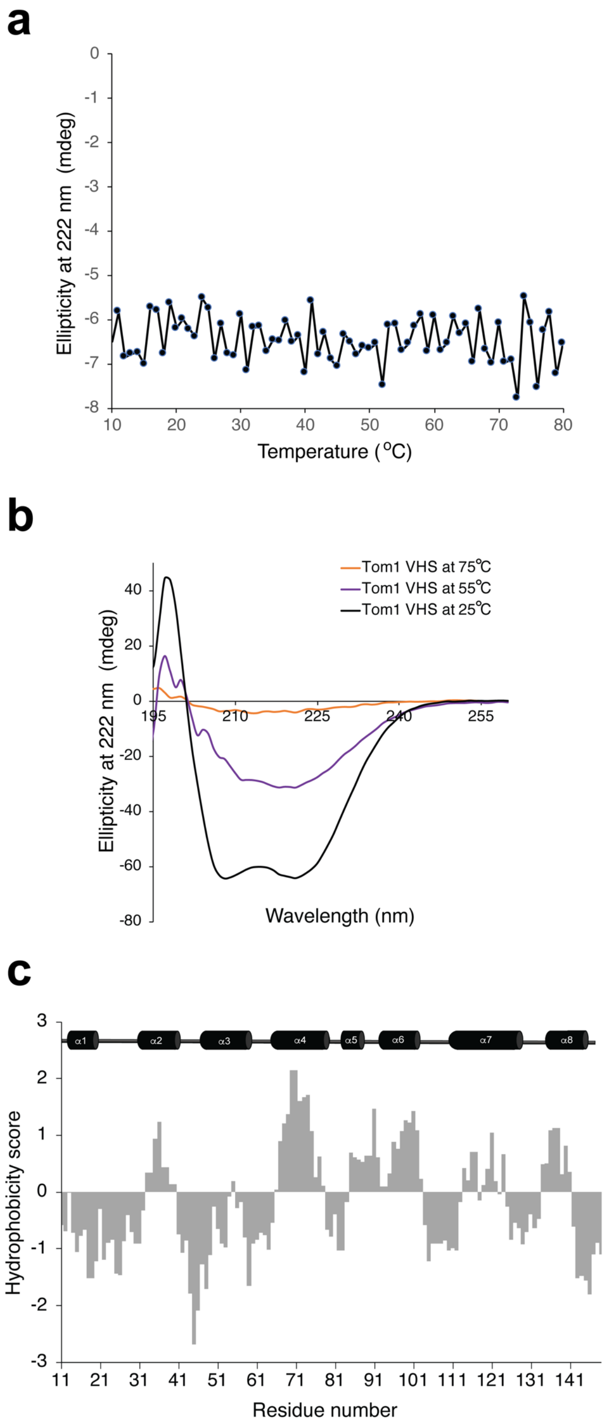


**Figure S1.** Thermal instability of the Tom1 VHS domain. (a) A refolding trial carried out by cooling the thermally denatured Tom1 VHS. (b) Far-UV CD spectra of Tom1 VHS at 25°C (blue), 55°C (purple), and 75°C (red). (c) Kyte and Doolittle hydrophobic score for the Tom1 VHS domain. The secondary structure of Tom1 VHS is shown at the top.

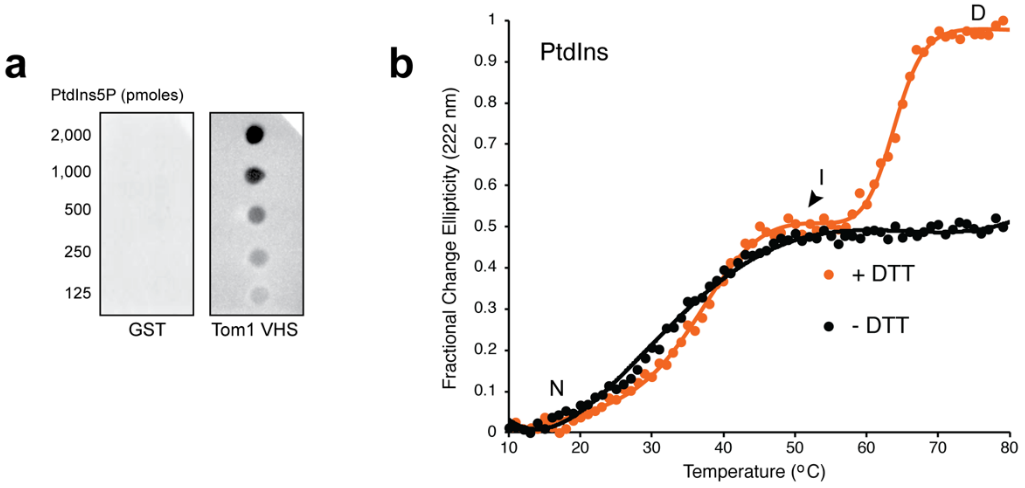


**Figure S2.** Effects of phospholipids on Tom1 VHS stability. (a) Lipid-protein overlay assay showing binding of Tom1 VHS to c16-PtdIns5P at the indicated amounts. (b) Fractional change in ellipticity at 222 nm with increasing temperature for Tom1 VHS pre-incubated with 12-fold of PtdIns without (black) and with DTT (orange).

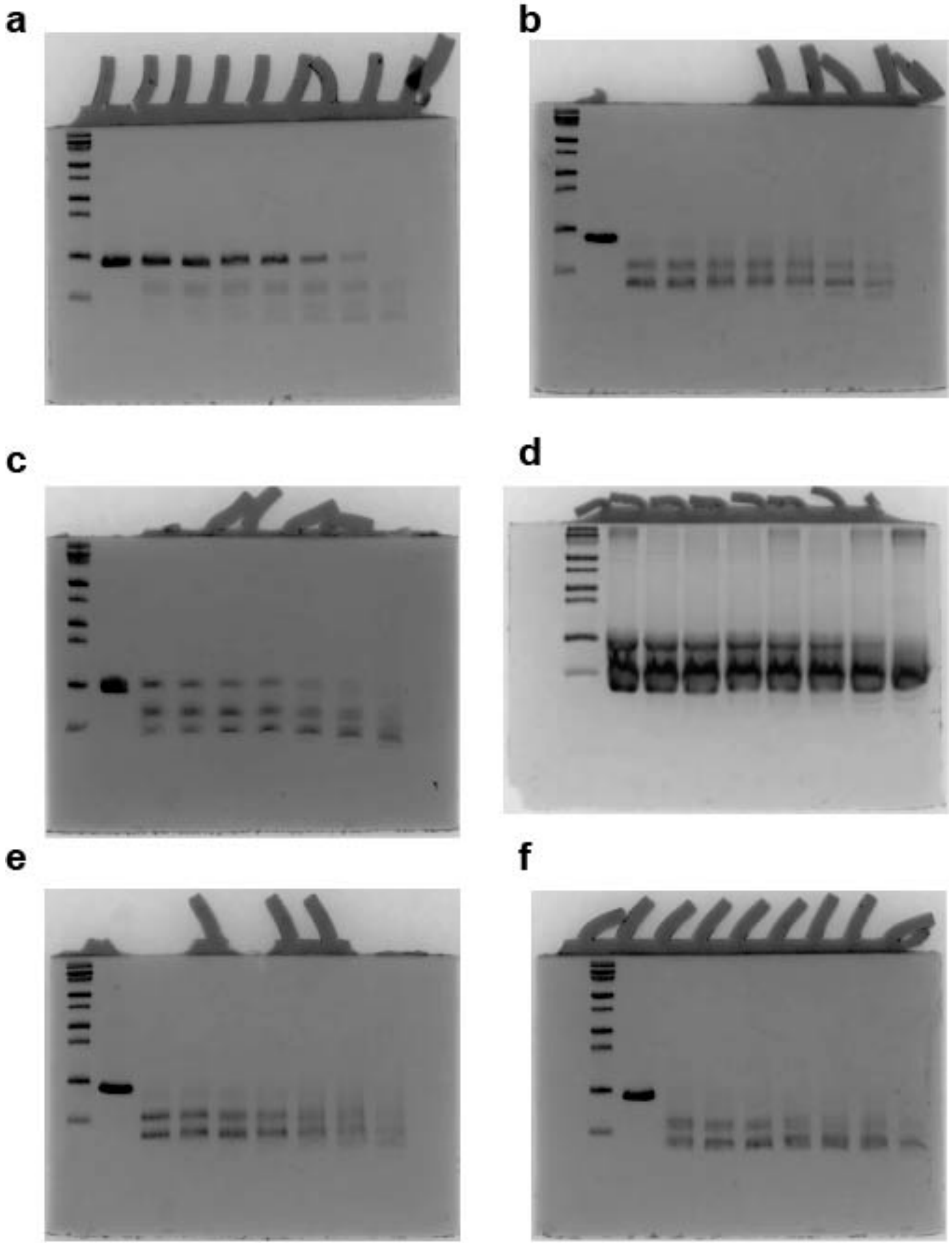


**Figure S3.** Trypsin limited proteolysis of Tom1 VHS domain in the absence and presence of ligands. Full SDS-PAGE scan images of trypsin-mediated proteolysis of Tom1 VHS in the absence (a) and presence of PtdIns5P (b), PtdCho (c), ubiquitin (d), PtdIns3P (e), and PtdIns (f).

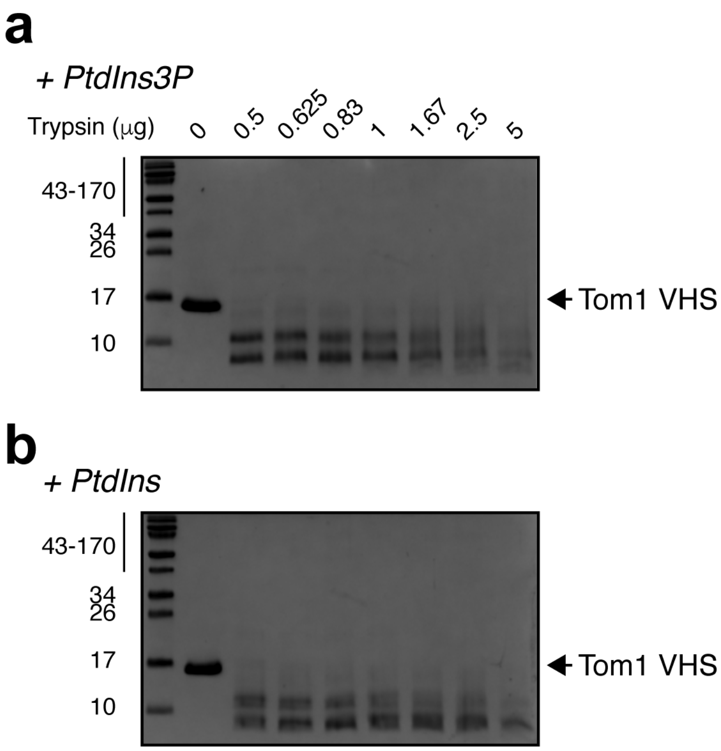


**Figure S4.** Lipid-dependent destabilization of the Tom1 VHS domain. SDS-PAGE showing the trypsin limited proteolysis of Tom1 VHS in the presence of 12-fold excess of PtdIns3P (a) and PtdIns (b).

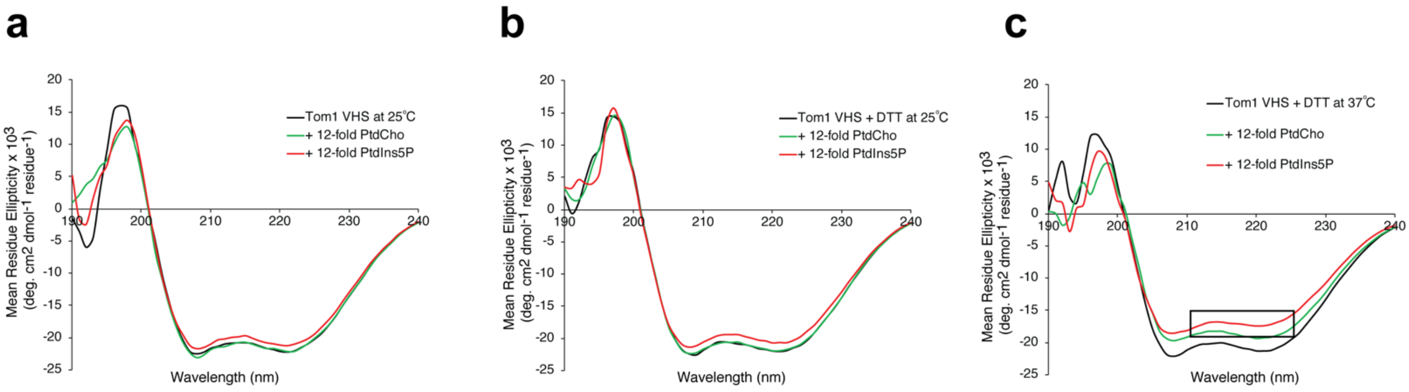


**Figure S5.** (a) Far-UV CD spectra of Tom1 VHS (20 μM) in the absence (black) and presence of 12-fold of PtdIns5P (red) or PtdCho (green) at 25°C. (b) Far-UV CD spectra of Tom1 VHS (20 μM) in 100 μM DTT in the absence (black) and presence of 12-fold PtdIns5P (red) or PtdCho (green) at 25°C. (c) Far-UV CD spectra of Tom1 VHS (20 μM) in 100 μM DTT in the absence (black) and presence of 12-fold PtdIns5P (red) or PtdCho (green) at 37°C. The loss of signal at 218-222 nm is boxed.

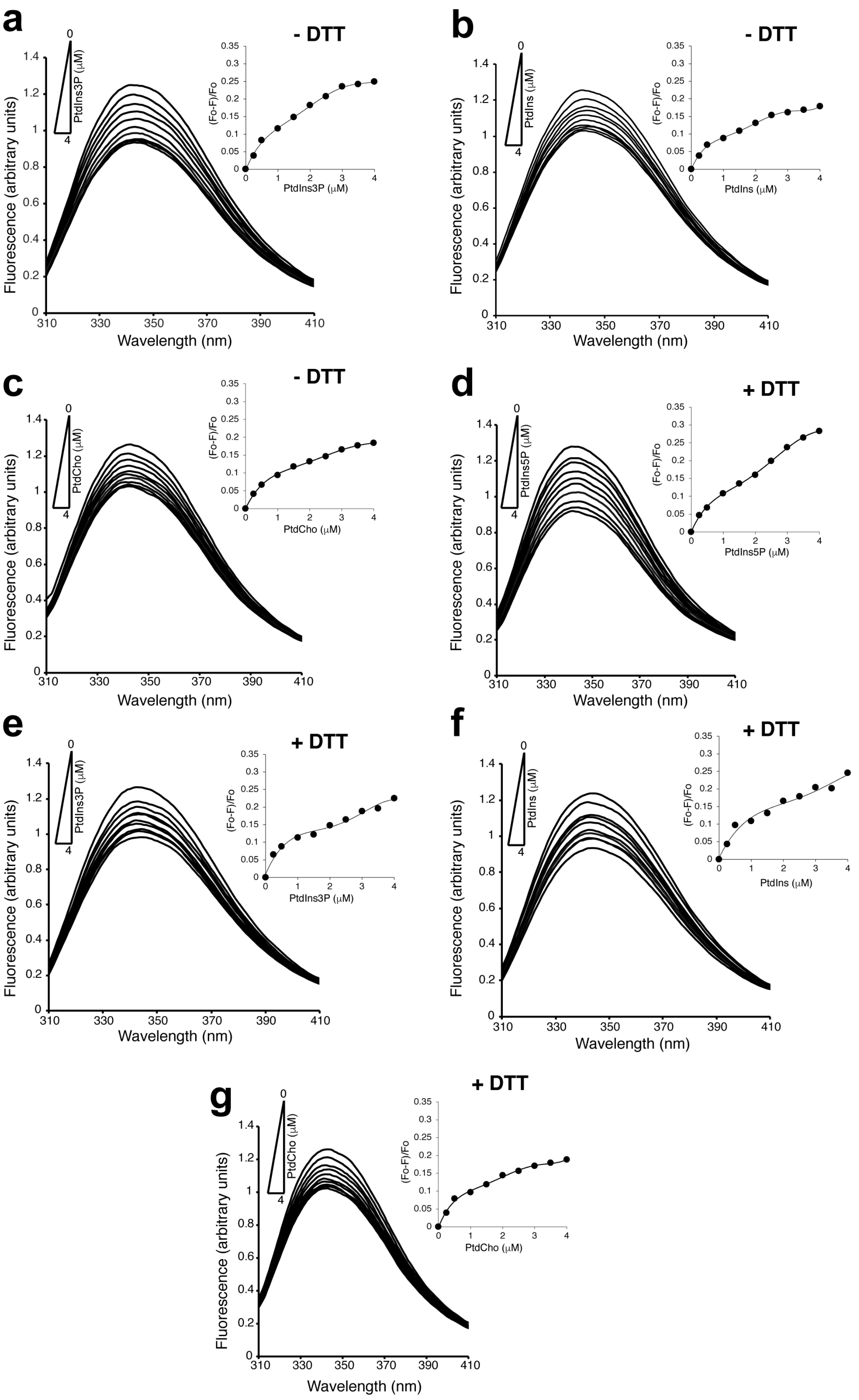
a
- DTT
PtdIns3P (μM)
0
4
Fluorescence (arbitrary units)
Wavelength (nm)
(Fo-F)/Fo
PtdIns3P (μM)
b
- DTT
PtdIns (μM)
Fluorescence (arbitrary units)
Wavelength (nm)
(Fo-F)/Fo
PtdIns (μM)
c
- DTT
PtdCho (μM)
Fluorescence (arbitrary units)
Wavelength (nm)
(Fo-F)/Fo
PtdCho (μM)
d
+ DTT
PtdIns5P (μM)
Fluorescence (arbitrary units)
Wavelength (nm)
(Fo-F)/Fo
PtdIns5P (μM)
e
+ DTT
PtdIns3P (μM)
Fluorescence (arbitrary units)
Wavelength (nm)
(Fo-F)/Fo
PtdIns3P (μM)
f
+ DTT
PtdIns (μM)
Fluorescence (arbitrary units)
Wavelength (nm)
(Fo-F)/Fo
PtdIns (μM)
g
+ DTT
PtdCho (μM)
Fluorescence (arbitrary units)
Wavelength (nm)
(Fo-F)/Fo
PtdCho (μM)

**Figure S6.** Changes in tryptophan fluorescence emission of Tom1 VHS induced by phospholipids. (a-c) Tom1 VHS with increasing concentrations of PtdIns3P (a), PtdIns (b), and PtdCho (c) in the absence of DTT. (d-g) Tom1 VHS with increasing concentrations of PtdIns5P (d), PtdIns3P (e), PtdIns (f), and PtdCho (g) in the presence of 1.25 μM DTT. Insets: Extent of phospholipid-dependent quenching of Tom1 VHS fluorescence signal.
